# Prediction of Epileptic Seizures based on Mean Phase Coherence

**DOI:** 10.1101/212563

**Authors:** Gorish Aggarwal, Tapan K. Gandhi

## Abstract

Around 1-2 % of the world's population suffers from epilepsy and 20 % of that vulnerable mass can't be cured through surgery or medicine. Epileptic seizures often occur unpredictably and may cause serious damage to the patient in adverse situations, for e.g. getting a seizure while driving or crossing a road can be fatal. In such a scenario, a reliable mechanism to predict the onset of seizure beforehand is much desirable. In this study, A reliable real-time technique for prediction of epileptic seizures is presented. Mean Phase Coherence (MPC), as a measure of phase synchronization, is used to predict impending seizures in a multi-channel EEG data. It was found that during the pre-ictal stages, MPC values and thus phase synchronization between various channels was found to drop significantly below the level in a non-ictal EEG signal. The range of the prediction horizon for seizures varied from 4-10 mins and prediction of impending seizure attack is possible for 8 out of 10 test cases.

## I. INTRODUCTION

Epilepsy is a chronic brain disorder characterised by recurrent seizures [1], [2]. They are brief stretches of involuntary movement affecting either a part of the body or the entire body and occur due to the excessive electrical discharges in a group of brain cells. Seizures can vary in their origin location, distance of spread as well as the time and frequency of occurrence.

According to World Health Organization, Epilepsy affects around 50 million people worldwide, making it one of the most common neurological disease, along with Stroke and Alzheimer's disease [3]. The lack of knowledge combined with non-existence of proper diagnosis and treatment methods for epilepsy makes this problem enormous for people of developing countries.

There are no known causes associated with epilepsy in most of the cases. However sometimes, brain injury, stroke and developmental abilities have been identified to cause epilepsy. The major symptom is the continued occurrence of seizure attacks. Since these attacks are highly unpredictable in nature and magnitude, they can cause serious damage to the patient in adverse situations. Thus, a low cost, easily accessible, point of care testing method for the detection and prediction of seizure is required, to monitor the patient's mental condition and warn him in advance of an impending seizure attack.

## II. LITERATURE REVIEW

Expert models have been built for automated detection of seizures from EEG signature with high accuracy [4], [5]. Techniques such as Neural Networks [6] and k-NN classifier [7] are easily able to classify EEG signatures to ictal or nonictal states. However, prediction of epileptic seizure is a relatively new field as compared to their detection. Earlier it was believed that the occurrence of seizures in brain is purely random and its start is also abrupt. However, in the late 1990s, a hypothesis [8] was proposed that the seizure starts building minutes to hours before the actual attack. This led to an interest and subsequent work on the prediction of seizure. However, till now, a robust technique has not been developed to predict seizure in all situations.

Mormann et al. [9] showed that phase synchronization between different EEG channels could be used as a measure for detecting the pre-ictal stage. He used correlation coefficient as a statistical measure for phase synchronization between the EEG channels. In another paper, Mormann et al. [10] also states that both the linear and non-linear techniques show similar performance for the prediction of seizures, thereby indicating that changes in dynamics were not necessarily caused solely by non-linearity of features. Le Van Quyen et al. [11] used dynamical similarity of EEG recording with a reference state as a measure for the prediction of seizures.

Zheng et al. [12] used a Bi-variate Empirical Mode Decomposition (BEMD) to first decompose windows of each pair of EEG to intrinsic mode functions (IMF). BEMD was used due to the non-stationary nature of EEG signals. He observed that there was both increase and decrease in phase synchronization before the seizure onset. Shufang et al. [13] used spike rate to distinguish the pre-ictal stage from ictal stage. It is based on the hypothesis that the spike rate increases as one moves from the inter-ictal state to pre-ictal state. They were able to predict an oncoming seizure with a sensitivity of 75.8 % and false prediction rate of 0.09/hr. Teixeira et al. [14] used multi-class SVM classifier and univariate features to classify the EEG signal of a person into 4 states: inter-ictal, pre-ictal, ictal and post-ictal.

### III. DATA

The data sets used for prediction of seizure has been obtained from 10 Epileptic patients and 2 Normal Individuals. The EEG data was received for 18 brain electrode channels, 9 in each hemisphere. The recordings were sampled at a frequency of 200 Hz. 7 patients had cases of generalized epilepsy and remaining 3 had focal epilepsies. The data was obtained from the Grass Telefactor Twin3 EEG machine at the Neurology and Sleep Centre, Hauz Khas, New Delhi.

The EEG data of an epileptic patient which contains seizures can be classified into 4 major regions:

1. Inter-ictal: Duration in-between the seizures and at sufficient gaps (~3 hours) from any seizure activities.
2. Pre-ictal: Duration before and near to a seizure activities. Generally chosen as 1-hour before a seizure. This is the region in which our system should be able to predict that a seizure can occur in the near future.
3. Ictal: The duration between seizure beginning till its end.
4. Post-ictal: Duration just after the seizure end till the EEG signal goes back to normal.

The ictal stage as well as the specific electrodes containing these ictal patterns have been marked by an expert. These markings are taken as ground truth for estimating the pre-ictal period as well as prediction horizon.

## IV. METHODOLOGY

Mean-Phase Coherence is used as a parameter to estimate an impending seizure activity. Mean-phase coherence values were found to decrease before the onset of an epileptic seizure. It is a measure for phase synchronization between the signals. Phase Synchronization is defined as the locking of the phases of 2 oscillatory system [51]. It is calculated as:

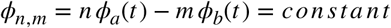

Instantaneous phase of the signal can be calculated using

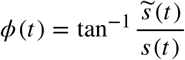

where 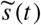 is the Hilbert transform of the signal given by

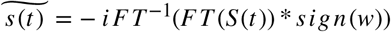

Here *FT* and *FT*^−1^ denotes the Fourier and the inverse Fourier transform respectively of the signal.

To measure the relative phase between these two signals a and b and to confine the phase between (−π/2*,π*/2), the following formula is used with values of *m* = *n* = 1

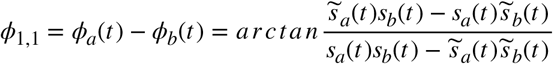

The mean phase coherence is calculated as:

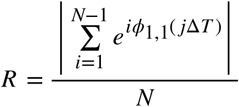

Studies show that the mean phase coherence [5] as a measure of phase synchronization between the EEG channels changes significantly during the pre-ictal stage when the brain waves shows signs of seizure formation as compared to the inter-ictal stage. This change in phase synchronization was found to occur before the actual seizure of seizure. Therefore, by measuring the value of mean phase coherence between channels, the onset of seizures can be predicted before their actual occurrence. The following steps were used:

### A. Pre-processing of data

The initial EEG data from 20 channels was filtered with a band pass filter from 0.5 Hz to 40 Hz and a notch filter at 50 Hz (to remove interference from the power supply of 50 Hz). The data sampled at frequency of 200 Hz was divided into windows of 5 seconds each.

### B. Calculation of Mean phase coherence (MPC)

The MPC values were calculated for each of the windows between 2 pairs of electrodes. The electrodes were chosen such only the coherence values between the neighbouring pairs of electrodes in each hemisphere were observed. This reduced the number of pairs to 24 from 153 (for 9 electrodes in each hemisphere). The above step helps in reducing the computational complexity as well as removes the redundancy in the data. The Hilbert transform and MPC were then calculated using the above equations.

### C. Thresholding and Classification

By thresholding the running sum of areas between the MPC values and their mean, it can be found whether a seizure attack is imminent in the near future. The threshold value used in our case for 10 windows i.e. 50 seconds was −0.4. Whenever the area between the graphs for the last 10 windows went below this value, the alarm would be turned on to warn the patient of an impending seizure. This threshold value was found to work fairly well for all the cases.

## V. RESULTS

In this part, results for seizure prediction using mean phase coherence as a parameter have been presented. The results are separated for different scenarios: with focal epilepsy, generalized epilepsy and a normal case. Mean phase coherence values were found to decrease significantly during the pre-ictal and rise again during the ictal stages. For a patient with focal epilepsy, only signal from electrodes placed around the focal point of seizures (available from patient history) showed a decrease in mean phase coherence whereas for a generalized seizure, the effect was visible in most of the electrodes. In the individual with no epilepsy, no changes in MPC during the entire duration of EEG recording were observed.

The results of each case have been discussed separately:

### A. Patient with Focal Epilepsy

This patient had a history of focal epilepsy in the occipital lobe. In the recorded EEG data for the patient, the seizure starts around the time 00:13:01 i.e. 156th window for a window size of 5 seconds. Fig 1 shows the EEG signature for this patient at a particular seizure instant.

**Figure 1:**
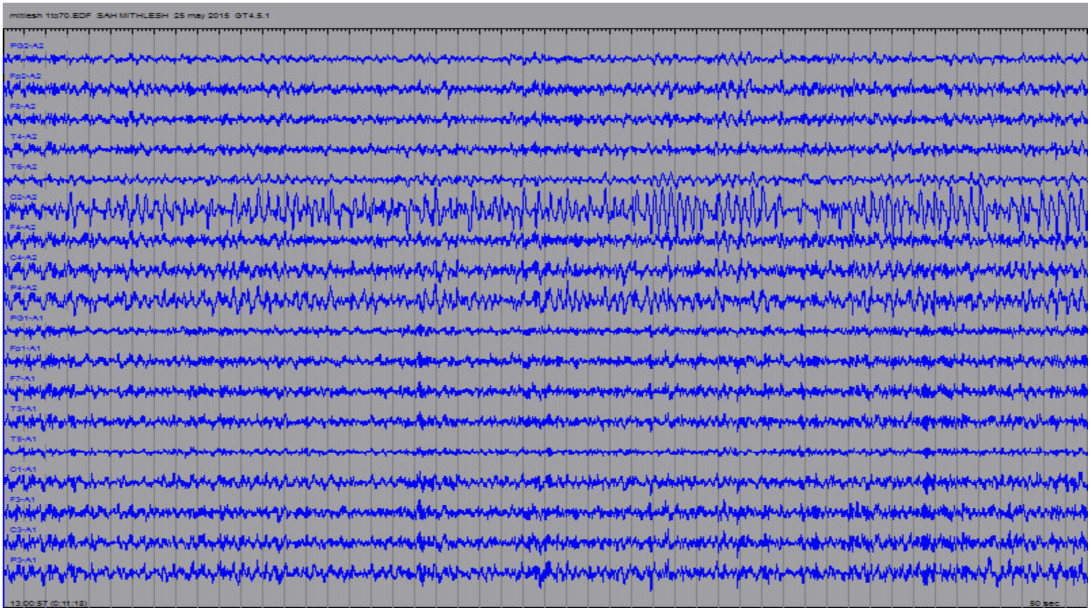
EEG signature for Patient 1 with focal epilepsy.

Fig 2 shows the value of MPC between the channels O2 and T6 for all the windows. The red line corresponds to the running mean values of the MPC and the green line shows the smoothed values of the graph, smoothing done by a moving average filter of length 4. Before the seizure actually occurs at 156th window filter of length 4. Before the seizure actually occurs at 156th window, it can be seen that the MPC values goes below the mean for a long duration before finally rising back up at the seizure time. Though there is variation of MPC throughout the recording period, the drop below the mean just before the seizure is the longest and most widely pronounced.

**Figure 2:**
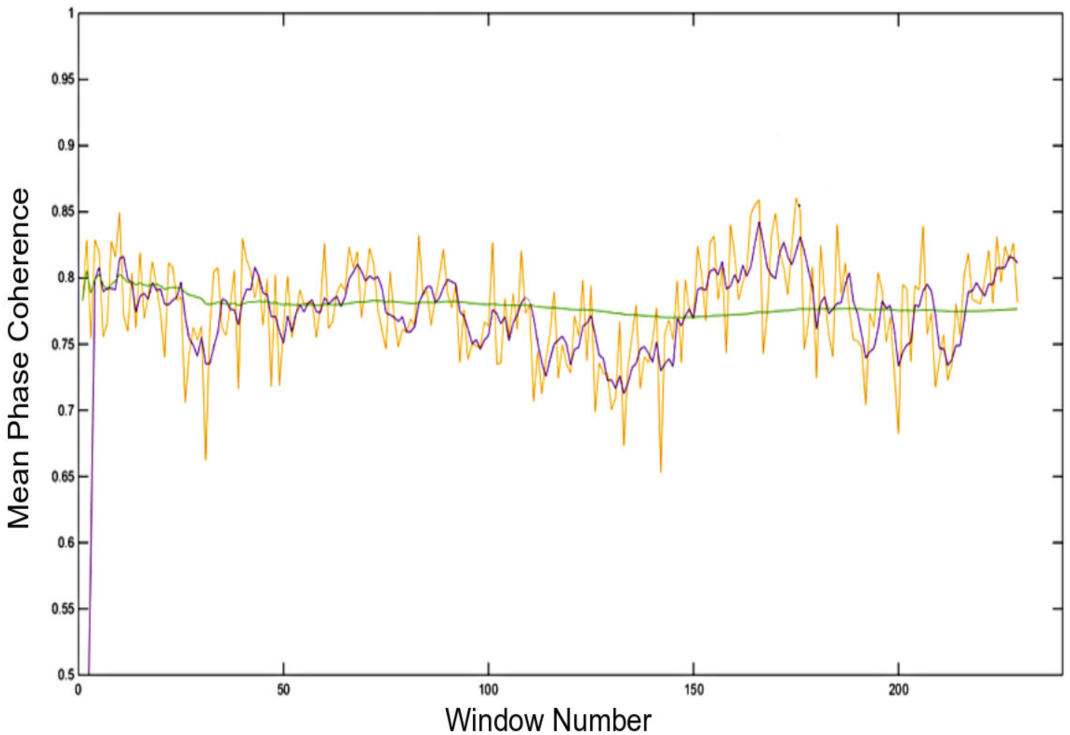
MPC for patient 1 for channels 02 and T6. Yellow: MPC, Violet: Smoothened MPC, Green: Running Mean of MPC

MPC values drop below the running mean around 93rd window and continues to remain below it for a long time, thus ensuring the fact that it is a pre-ictal stage and a seizure attack is on the horizon. Prediction horizon for this case turned out to be around 5 minutes.

Fig 3 also shows the MPC values between O2 and P4. It can be seen that the drop below the mean value is also pronounced highly just before the seizure.

**Figure 3:**
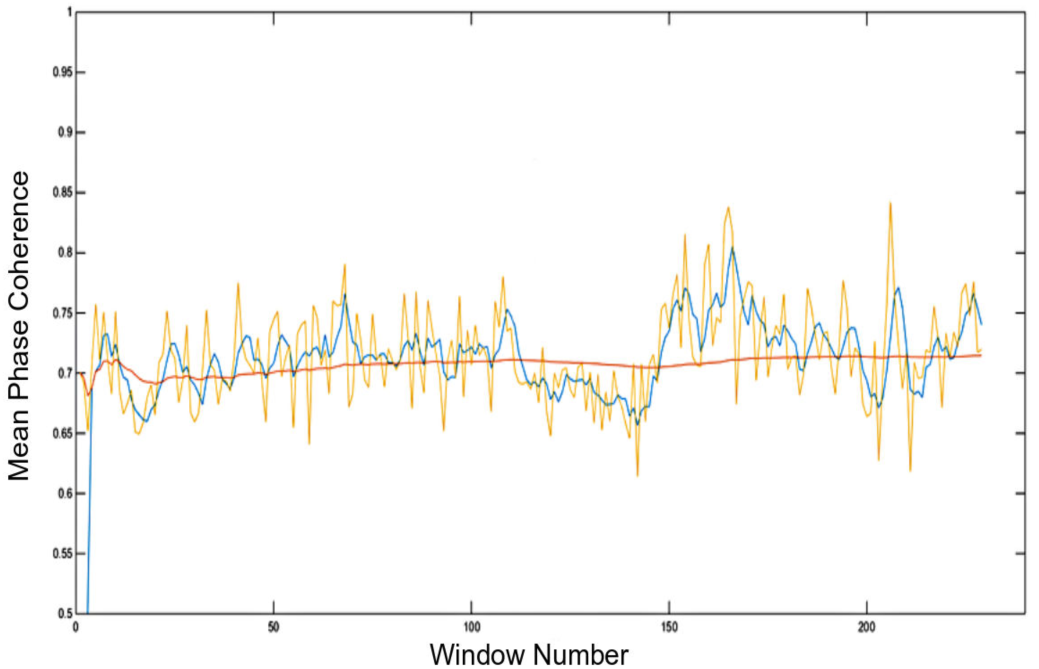
MPC for patient 1 for channels O2 and P4. Yellow: MPC, Blue: Smoothened MPC, Green: Running Mean of MPC

However, in Fig 4, for channels T4 and F8, there is no drop in MPC values, thus implying that the fact no seizure activity can be developed in these channels. Therefore, the impact of phase synchronization decrease in the pre-ictal period is not reflected in the entire brain in case of a focal epilepsy.

**Figure 4:**
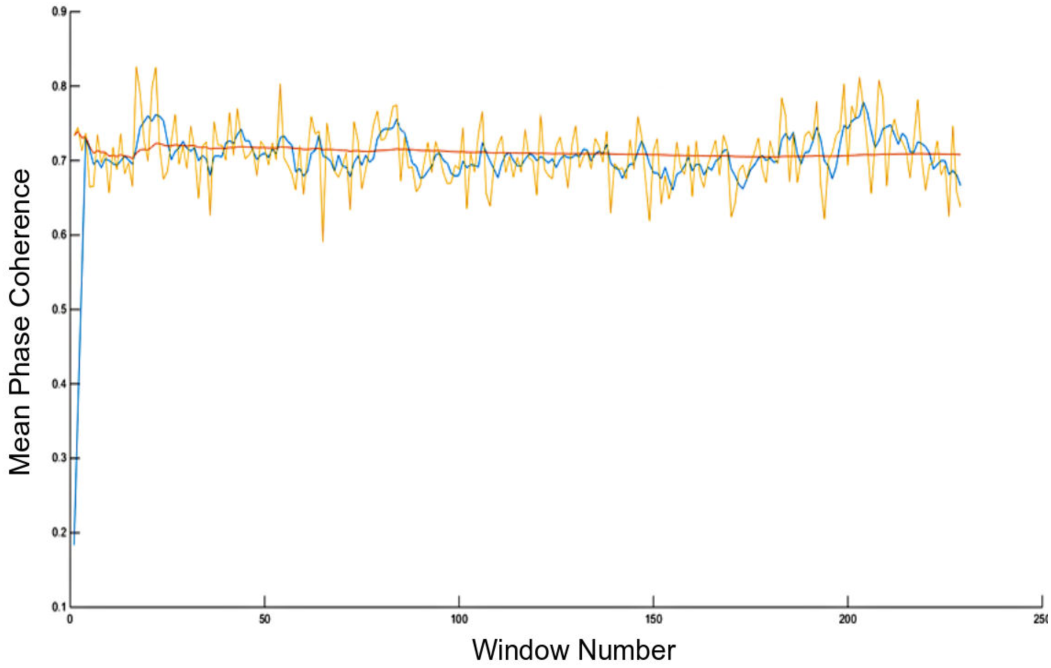
MPC for patient 1 for channels F8 and T4. Yellow: MPC, Blue: Smoothened MPC, Green: Running Mean of MPC

### B. Patient with Generalized Epilepsy

This patient has a history of generalized epilepsy in the brain. Fig 5 shows a snapshot of EEG during seizure activity for this patient. It can be seen that the EEG pattern shows large amplitude during the seizure and also very small gaps between 2 seizure duration. The current frame size of image is around 100 seconds. The seizure is starting at 00:06:10. The pre-ictal stage available for this patient is very short lasting for only around 5 minutes. Hence, the drop of the curve below the mean is not very distinctive. Fig 6 shows the filtered MPC values between the channels O2 and P4. From Fig. 6, for the region between the 10th window and the 75th window, there is a drop below the mean of the filtered MPC values. There is a similar drop observed for MPC values between channels in right hemisphere such as C3 and P3. Prediction horizon for this case is around 7 minutes.

**Figure 5:**
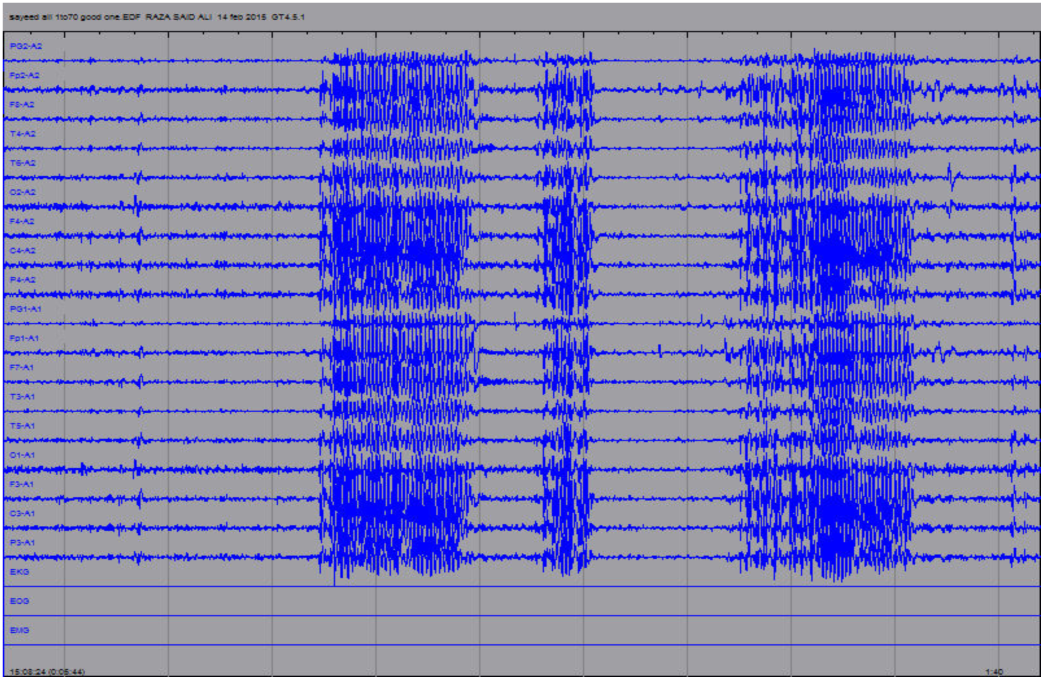
EEG signature for patient 2 with Generalized Seizure.

**Figure 6:**
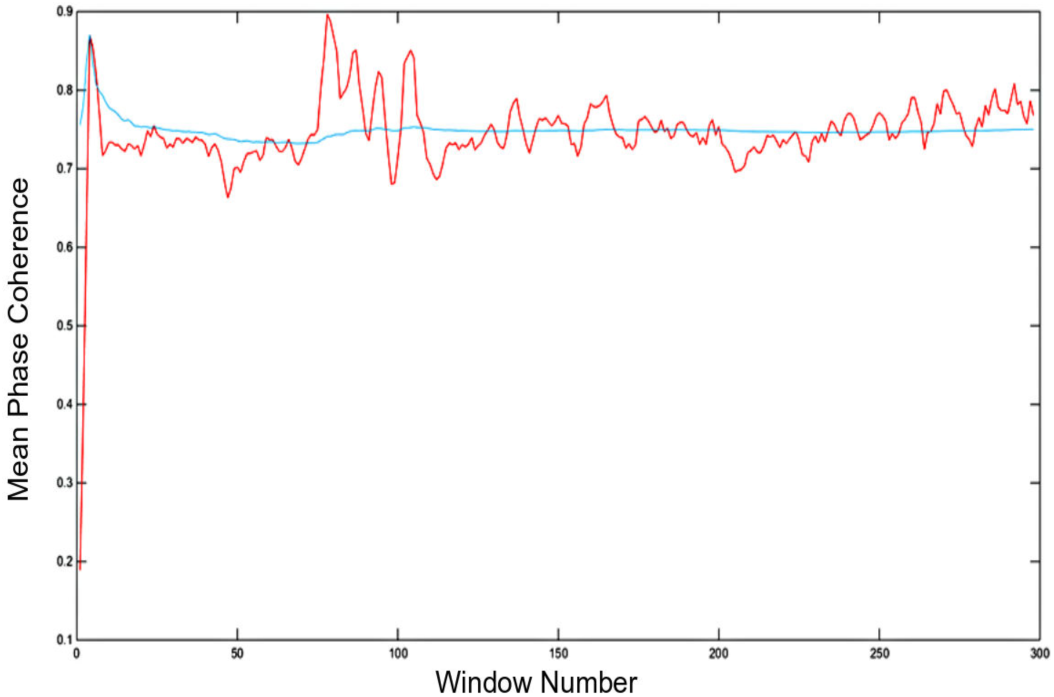
MPC for patient 2 for channels 02 and P4. Red: Smoothened MPC, Blue: Running Mean of MPC

### C. Normal Individual

The mean phase coherence between different channels were plotted for a normal person in asleep state. Fig 7 corresponds to the MPC between channel T4 and F8. From the graph, it can be seen that the mean phase coherence values neither increase, nor decrease considerably. This trend has been observed uniformly across most of the EEG channel pairs. There is no trend of MPC going below the running average, so there is no prediction of an impending seizure. This was confirmed by the ground truth values.

**Figure 7:**
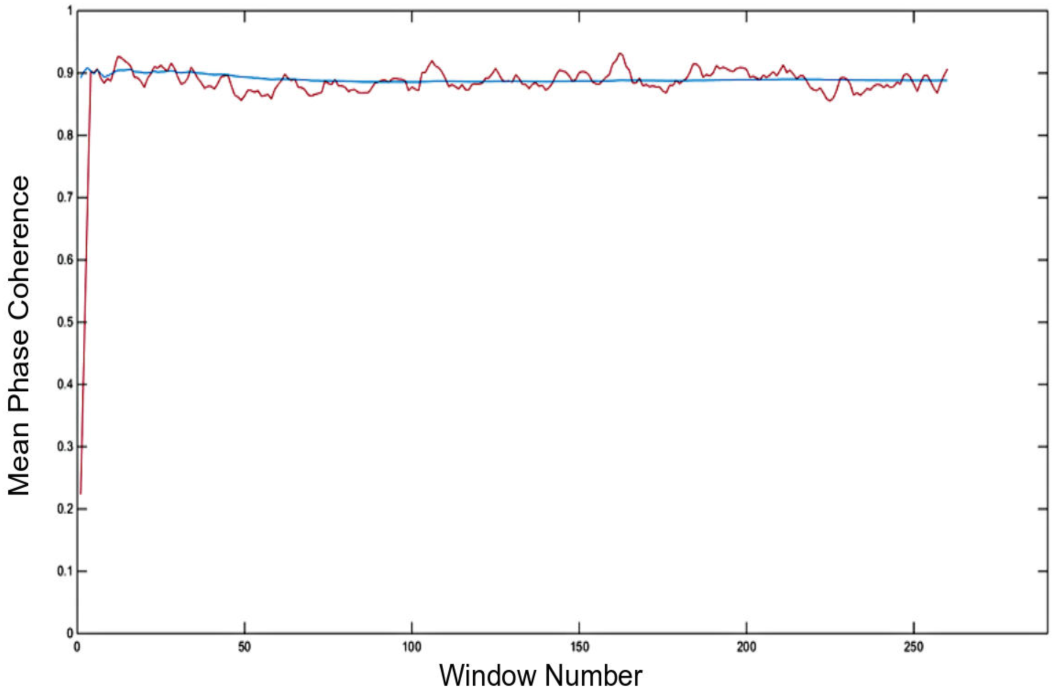
MPC for normal person for channels F8 and T4. Red: Smoothened MPC, Blue: Running Mean of MPC

## VI. CONCLUSION

The mean phase coherence values were found to decrease below a threshold value before a seizure was about to occur for 8 patients out of 10. Focal Epileptic seizures were predicted in all 3 cases while only 5 out of 7 cases of generalized epileptic seizure were predicted before time. The drop was observed in significant number of EEG channels (scattered in both hemispheres) for patients with generalized epilepsy whereas it was only seen in particular channels for patient with focal epilepsy. This is important as it shows the validity of use of MPC as a measure for phase synchronization of channels and to determine the onset of a seizure attack. The prediction horizon for all positive cases was between 4 - 10 mins before the actual onset of seizures. Both non-epileptic subjects did not show any significant change in MPC values for the entire duration of EEG recording. Thus, it was found that in the minutes leading to a seizure attack, there is drop in phase synchronization between different channels of EEG and this drop is specific to those electrode of brain regions where seizure activity is observed.

